# Functional antibodies exhibit light chain coherence

**DOI:** 10.1101/2022.04.23.489267

**Authors:** David B. Jaffe, Payam Shahi, Bruce A. Adams, Ashley M. Chrisman, Peter M. Finnegan, Nandhini Raman, Ariel E. Royall, FuNien Tsai, Thomas Vollbrecht, Daniel S. Reyes, Wyatt J. McDonnell

## Abstract

The vertebrate adaptive immune system modifies the genome of individual B cells to encode antibodies binding particular antigens^1^. In most mammals, antibodies are composed of a heavy and a light chain which are sequentially generated by recombination of V, D (for heavy chains), J, and C gene segments. Each chain contains three complementarity-determining regions (CDR1-3), contributing to antigen specificity. Certain heavy and light chains are preferred for particular antigens^2–21^. We considered pairs of B cells sharing the same heavy chain V gene and CDRH3 amino acid sequence and isolated from different donors, also known as public clonotypes^22,23^. We show that for naive antibodies (not yet adapted to antigens), the probability that they use the same light chain V gene is ∼10%, whereas for memory (functional) antibodies it is ∼80%. This property of functional antibodies is a phenomenon we call *light chain coherence*. We also observe it when similar heavy chains recur *within* a donor. Thus, though naive antibodies appear to recur by chance, the recurrence of functional antibodies reveals surprising constraint and determinism in the processes of V(D)J recombination and immune selection. For most functional antibodies, the heavy chain determines the light chain.

## A novel approach to grouping antibodies reveals light chain coherence in functional antibodies

A central challenge of immunology is to group antibodies by function. Ideally, antibodies in such groups would share both an epitope and complementary paratopes dictated by their protein sequences. Practically, small numbers of antibodies are assayed *in vitro* e.g., for functional activities such as neutralizing capability. Larger numbers of antibodies can be assayed for simple binding to a particular antigen. In the future, possibly, antibody properties might be understood at scale from sequence information alone, perhaps via structural modeling, and via that antibodies might be grouped. However, at present, in lieu of a sufficiently large dataset with multiple antigen specificities using cells from multiple humans or donors, it is impossible to assess the validity of *any* functional grouping scheme.

We can however make some inferences. All the antibodies within a clonotype—a group of antibodies that share a common ancestral cell which arose in a single donor—usually perform the same function. A clonotype can therefore be treated as the minimal functional group of antibodies. Next, as has been observed, nature repeats itself by creating similar clonotypes that appear to have the same function^**2-21**^, and these might be combined into groups. Such recurrences have been observed between donors, but also occur within single donors, as we will demonstrate. Regardless, such recurrences arise *after* recombination randomly creates a vast pool of potential antibodies; recurrences arise through *selection* from that pool.

Specific examples suggest that sequence similarity can guide the way to understanding functional groups. For example, in the case of influenza virus, antibodies binding the anchor epitope of the haemagglutinin stalk domain reuse four heavy chain V genes and two light chain V genes^21^. A similar observation has been made in the case of Zika virus, where a protective heavy-light pair *IGVH3-23*/*IGVK1-5* is observed in multiple humans, which also cross-reacts with dengue virus^16^. Even in the setting of human immunodeficiency virus infections, which lead to diverse and divergent viruses within a single human, recurrent and ultra-broad neutralizing antibodies such as the VRC01 lineage emerge, with subclass members using combinations of *IGHV1-2* and *IGHV1-46* heavy chains paired with *IGKV1-5, IGKV1-33, IGKV3-15*, and *IGKV3-20* light chains^24^.

Motivated by these examples, we set out to answer the following simple question—do *unrelated* B cells with similar heavy chains also have similar light chains? We exclude *related* cells (*i*.*e*., those in the same clonotype) because they use the same VDJ genes *by definition*.

We generated a large set of paired V(D)J data to investigate this question. Using peripheral blood samples from four unrelated humans (**Methods**), we captured and sequenced paired, full-length antibody sequences from a total of 1.6 million single B cells of four flow cytometry-defined phenotypes^25,26^: naive, unswitched memory, class-switched memory, and plasmablasts (**Methods, Extended Figure 1**). For each cell, we obtained nucleotide sequences spanning from the leader sequence of the V gene through enough of the constant region to determine the isotype and subclass of the antibody (**Methods**).

We computationally split these antibody sequences into two types: (1) naive and (2) memory. To do so, we inferred V gene alleles for each of the four donors (**Methods**), and then for each B cell used the inferred alleles to estimate the number of somatic hypermutations which occurred outside the junction regions, including both chains. We refer to this number as donor reference distance (*d*_*ref*_). We labeled an antibody sequence as **naive** if it had no mutations relative to the inferred germline (*d*_*ref*_ = 0), and as **memory** otherwise (*d*_*ref*_ > 0). We compared these categories to the flow sort categories (**Extended Table 1**), noting general consistency but also that computational sorting was somewhat more accurate. Approximately 80.0% of cells flow sorted as naive were naive by computational analysis, with a maximum accuracy of 90.9% for donor 2. Conversely, just 0.2% of cells flow sorted as memory were computationally naive. During library preparation we exhausted the supply of memory cells and deliberately mixed them in some libraries (e.g. switched B cells plus naive B cells) to best exploit capacity. Our computational sorting also enabled us to make the best use of all the data.

Next we investigated the question described above, if for unrelated B cells, similar heavy chains imply similar light chains. We explored this question separately in memory and naive cells by considering pairs of cells, either both memory or both naive. We only considered pairs of cells with the same heavy chain V gene and the same CDRH3 length, and whose cells came from different donors. We divided the pairs into eleven sets by their CDRH3 amino acid percent identity, rounded down to the nearest 10%. Then for each set we computed its <u>light chain coherence</u>: the percent of cell pairs in which the light chain gene names were identical. We consider light chain V gene paralogs with “D” in their name to be identical in this work (e.g., *IGKV1-17* and *IGKV1D-17*).

We show the results of this analysis in **Figure 1**. For memory B cells found in separate donors having the same heavy chain V genes and 100% CDRH3 amino acid identity (2,813 cells), we found **82%** coherence between their light chains, whereas light chain coherence in naive cells (754 cells) was only **10%**. This makes sense, as naive cells have generally not yet been selected for functionality or undergone somatic hypermutation during an immune response. This finding implies that for memory cells, which bear functional antibodies and typically are the products of thymic and peripheral selection, heavy chain coherence implies light chain coherence. We note that light chain coherence does not imply heavy chain coherence (**Extended Figure 2**). We also note that if we instead define naive (CD19^+^IgD^+^CD27^±^CD38^±^CD24^±^) and memory cells (**Methods**) by flow cytometry, we find 86% concordance between light chains for memory, and 16% for naive.

**Figure 1.**
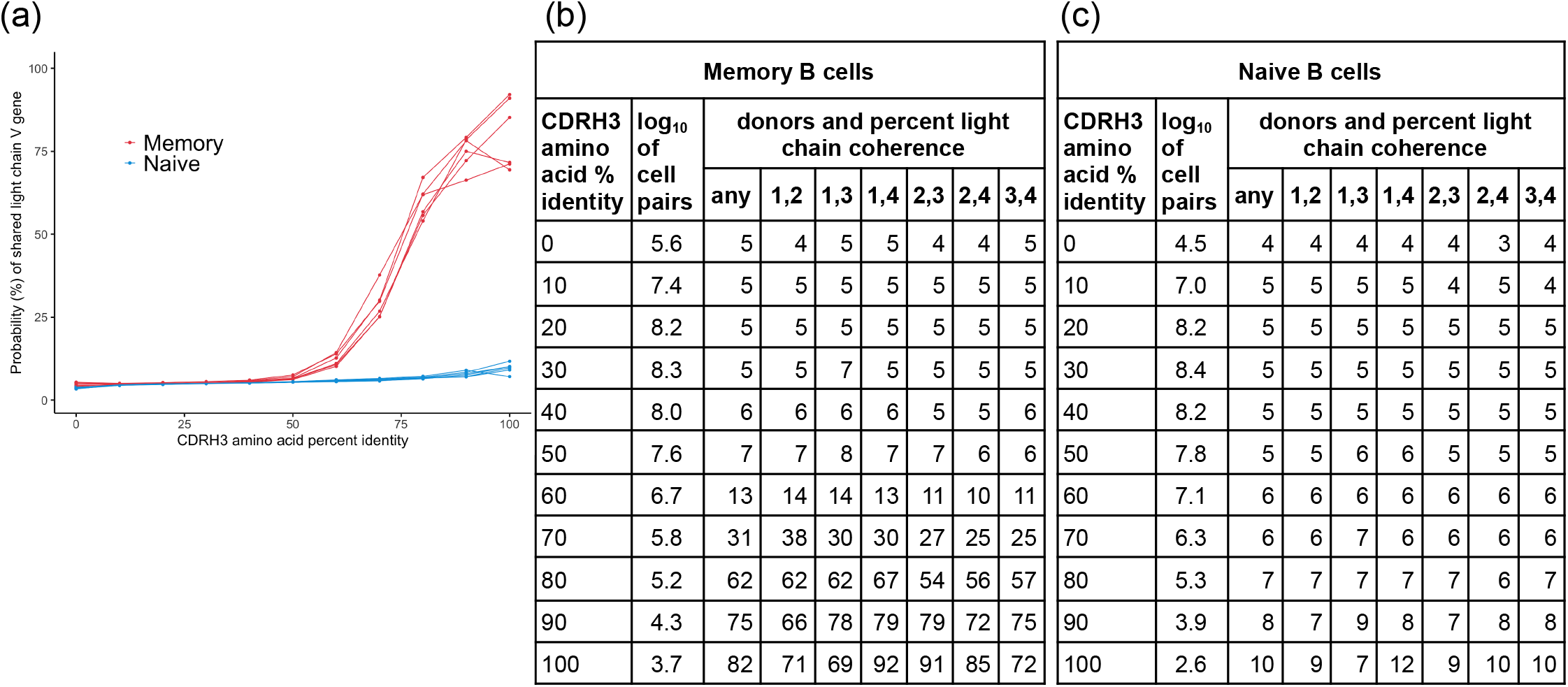
Light chain coherence is a property of public antibodies in memory B cells. **(a)**: Pairs of B cells were examined if (1) they had the same heavy chain V gene name, (2) they had the same CDRH3 length, (3) they were from different donors, and (4) both cells were either memory (red), or both were naive (blue), with one curve per pair of donors. The percent of cell pairs using the same light chain V gene (or paralog) is shown as a function of CDRH3 amino acid identity, rounded down to the nearest 10%. **(b**,**c)**: Light chain coherence values from the curves in (a) are displayed as tables. Light chain coherence varied depending on which donors’ antibodies were compared, as shown for particular pairs of donors 1, 2, 3, and 4.

Light chain coherence is still visible even if light chain V gene paralogs are not treated as the same gene, with light chain coherence of 64% for memory cells (**Extended Figure 3**). A more sophisticated approach might make *more* identifications, as sufficiently similar V genes should be functionally indistinguishable. In fact, light chain coherence can be observed without reference to genes at all. Given pairs of cells from different donors, we can compute their heavy and light chain edit distances. In that case, if the cells have the same CDRH3 amino acid sequence, then **78%** of the time, their light chain edit distance is ≤ 20, whereas without the CDRH3 restriction that is true only **9%** of the time (**Extended Figure 4**).

We also reasoned that light chain coherence could have nothing to do *per se* with multiple donors and could instead be investigated within a single donor. However, identifying genuine recurrence can be precarious. In principle one could base such an analysis on pairs of computed clonotypes, though an obvious concern would be that two computed clonotypes were in fact part of a single true clonotype that was incompletely grouped (*cf*. **Figure 3(c)**).

Therefore we restricted our attention to computed clonotypes that we were certain were truly separate. To do this we considered pairs of cells, each from the *same* donor, with *different* heavy chain gene names and equal CDRH3 lengths. These results are shown in **Figure 2**.

**Figure 2.**
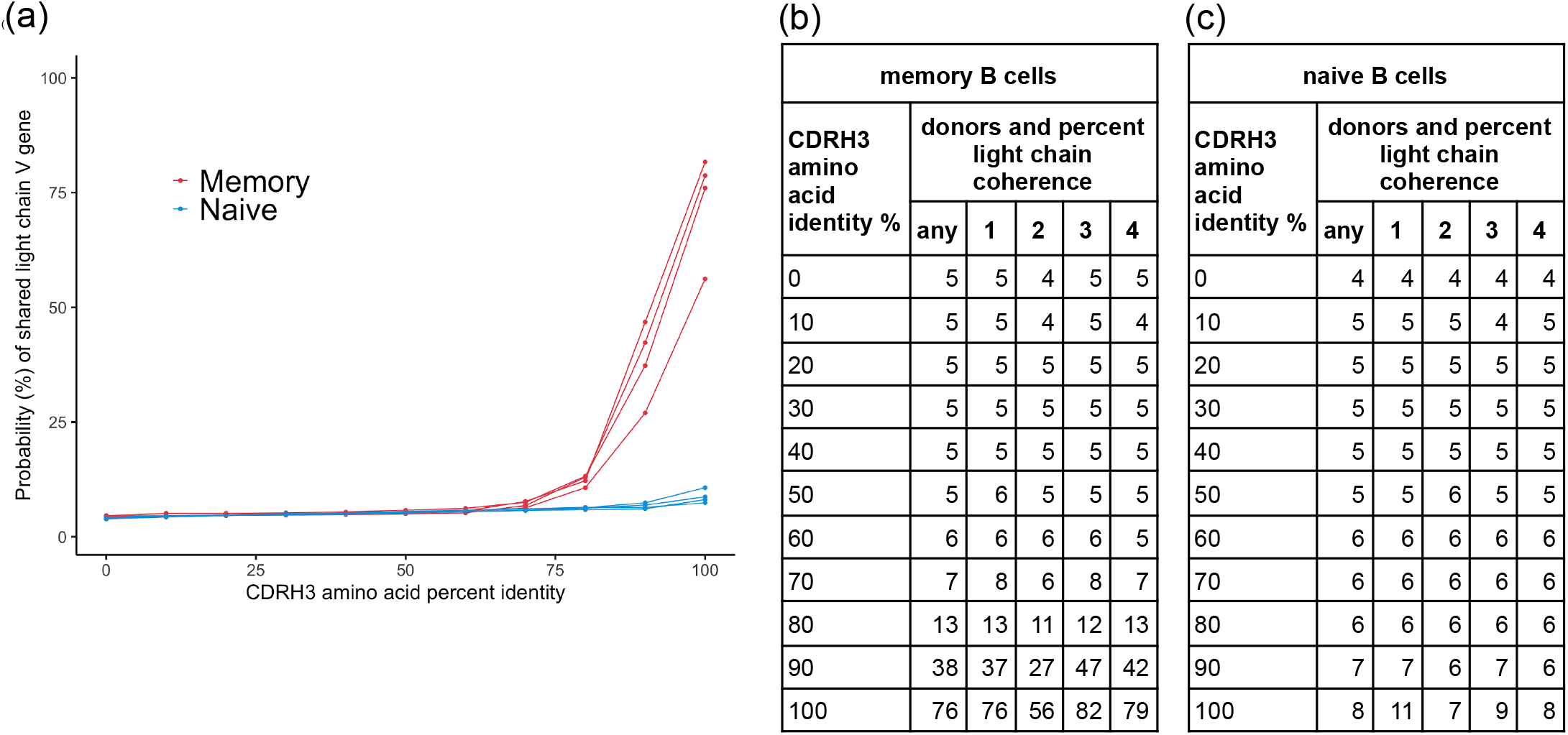
Light chain coherence is a property of private antibodies in memory B cells. Pairs of single B cells within a donor were examined if (1) they had the same CDRH3 length, (2) they used *different* heavy chain V genes, and (3) either both were memory (red), or both were naive (blue), with one curve for each donor shown in **(a)**. The percent of cells using the same light chain V gene (or paralog) is shown as a function of CDRH3 amino acid identity, rounded down to the nearest 10%. In **(bc)**, data for single donors 1, 2, 3, 4 are shown to exhibit dependence on them.

**Figure 3.**
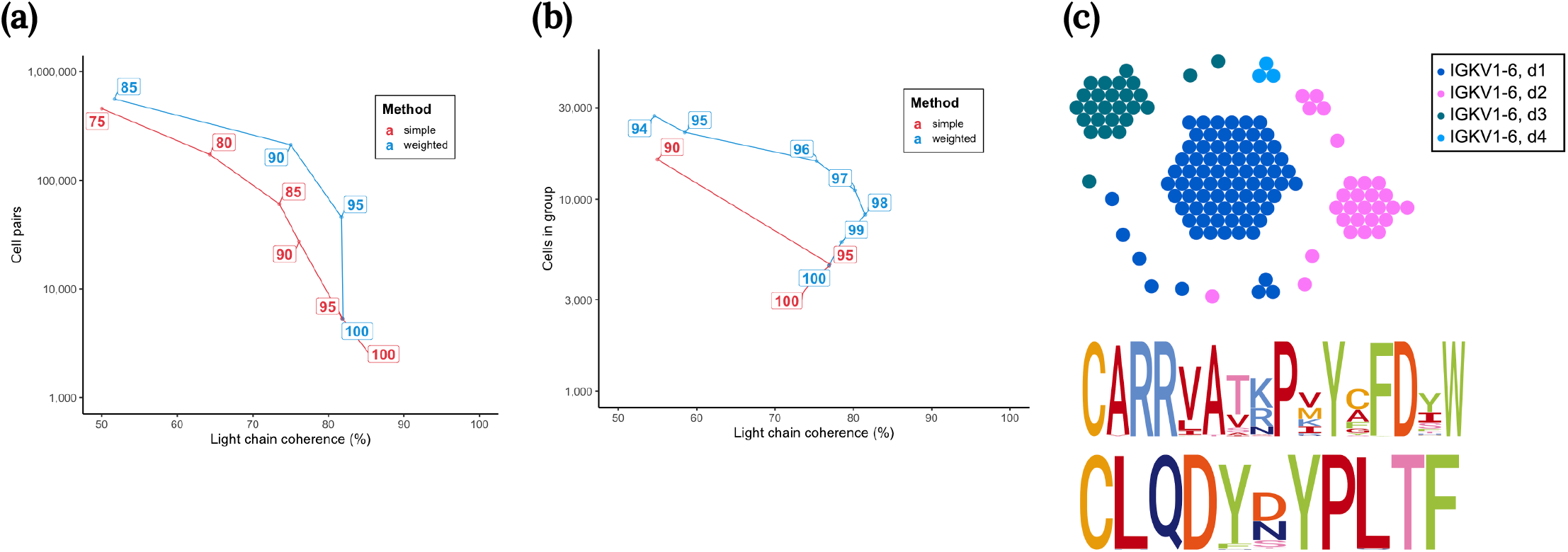
Substitution matrix optimization and transitive grouping of clonotypes. **(a)** Consider pairs of memory cells in the data, with each cell in a pair from a different donor, and with both cells sharing the same heavy chain V gene name and CDRH3 length. Red: for a given percent identity (points labeled by such), find all cell pairs satisfying the given percent identity on CDRH3 amino acids. Coordinates are given by the light chain coherence for these cells and the number of pairs. Turquoise: the same, but now use the weighted percent identity defined by the COSUM matrix M (see text). **(b)** Transitively group clonotypes, using a given percent identity or weighted percent identity. Now compare light chain coherence to the number of cells appearing in the groups. **(c) [top]** Using weighted percent identity 95%, the group for CDRH3 = CARRVATKPVYCFDYW is displayed. These cells use the heavy chain gene *IGHV4-59*. Each dot is a cell, and each cluster is a computed clonotype. All computed clonotypes use the light chain gene *IGKV1-6*, and all four donors (d1, d2, d3, d4) are present. The following J gene usage is observed: *IGHJ4*/*IGKJ1* or *IGHJ1*/*IGKJ1* or *IGHJ4*/*IGKJ4* (d1); *IGHJ4*/*IGKJ4* or *IGHJ3*/*IGKJ4* (d2); *IGHJ3*/*IGKJ3* (d3); *IGHJ3*/*IGKJ4* (d4), thus making it likely that at least 7 true clonotypes are present, but some of the 18 computed clonotypes might be merged in the true clonotypes. **[middle]** logo plot for CDRH3 amino acids. **[bottom]** logo plot for CDRL3 amino acids.

For memory cells, at 100% CDRH3 amino acid coherence, we saw **76%** light chain coherence, which is slightly lower than shown in **Figure 1** for public antibodies. This is unsurprising given that the condition on heavy chain coherence was relaxed. We would expect the true rates of coherence to be approximately equal. Thus, we provide evidence that light chain coherence appears to be a general property of antibodies with a common function.

So far, we have considered functional grouping of antibodies based on CDRH3 amino acid percent identity, as have others^15,17,19^. However, such grouping treats all amino acids as equal and cannot be optimal. Therefore, we used light chain coherence to develop an amino acid substitution matrix that might better reflect functional differences between particular amino acids in antibody sequences. To do this, for a given amino acid substitution matrix M, we defined the *weighted percent identity* of two amino acid sequences X and Y of length *n* with the following formula:

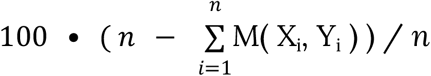

For the simple matrix having entry 0 for identical amino acids, and 1 otherwise, this is the same as percent identity. We started with this matrix and then partially optimized it for light chain coherence (**Methods**). This light chain coherence optimized substitution matrix (COSUM) M is exhibited as **Extended Figure 5**. As expected, highly similar amino acids tend to have low substitution penalties, e.g. M(L, I) = 0.1, whereas dissimilar ones have high values, e.g. M(L, P) = 8.0. In **Figure 3(a)**, we see how weighted percent identity using COSUM compares to ordinary percent identity. Briefly, for the same light chain coherence, use of COSUM-weighted percent identity is several times more likely to find that two antibodies are proximate. We *hypothesize* that light chain coherence is a proxy for actual functional coherence.

After considering pairs of cells, we turned our attention to transitive grouping of computed clonotypes. To understand what *transitive* means in this context, suppose that all clonotypes are to be placed in non-overlapping groups. A rule for this might be that two clonotypes go in the same group if they are similar, by some given criteria. However, this forces certain clonotypes to go in the same group even though they are *not* similar. Thus if X is similar to Y, they both go in the same group, and if Y is similar to Z, they both go in the same group, and therefore Y and Z must go in the same group, even if they are not similar. The two hops X to Y to Z define a transitive connection.

We considered only computed clonotypes consisting entirely of memory cells. As before, we grouped computed clonotypes (and therefore cells) together according to an identity requirement. Clonotypes were placed together in a group if one cell from one clonotype was found having the given identity with some cell in the other clonotype. This was done transitively. The process yielded groups of clonotypes (and therefore groups of cells), which could then be analyzed for light chain coherence by examining pairs of cells from the same group but from different donors.

As can be seen by examining comparing **(a)** and **(b)** of **Figure 3**, transitive grouping lowers coherence. This is because of the multiple “hops” involved in transitivity. In **Figure 3(c)**, a group computed using 95% weighted percent identity is exhibited. The light chain coherence for this group is 100%, and at least 7 and possibly up to 18 true clonotypes are present, representing independent recombination events (recurrences). Both heavy and light CDR3 sequences exhibit strong conservation.

## Observed public antibodies arise from low complexity V(D)J recombination

Recurrent antibodies observed in a small number of donors (such as we have) would be expected to arise from relatively common recombination events. We investigated this (**Figure 4**) by further exploring three questions.

**Figure 4.**
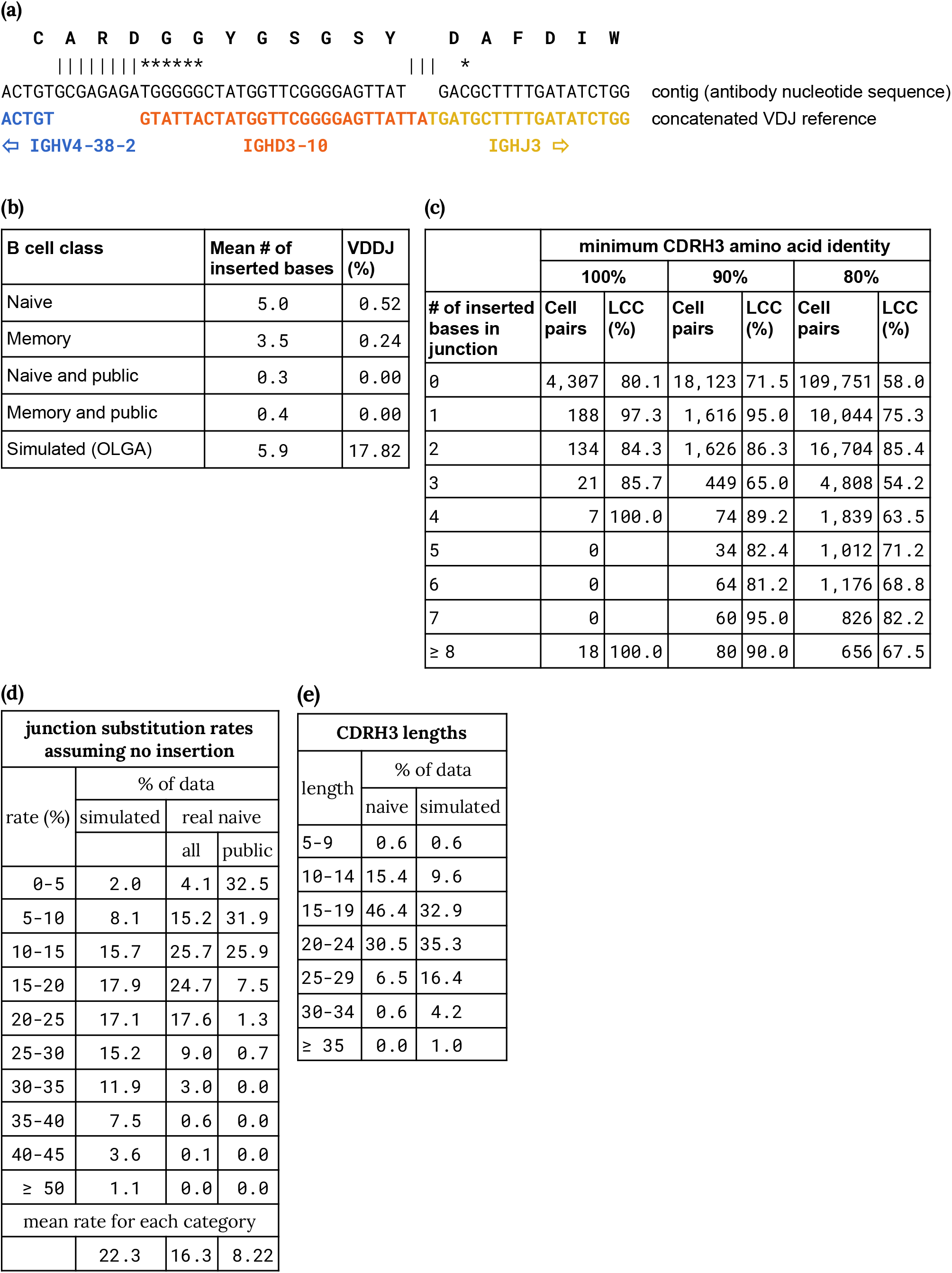
Heavy chain junction properties. Heavy chain junction sequences were aligned to concatenated reference sequences, allowing for VJ or VDJ or VDDJ, with up to two different D genes, and the most likely reference selected. We determined the number of bases inserted in the junction, relative to this reference (counting deletions separately). **(a)** The heavy chain junction region for a memory cell with the heavy chain junction CARDGGYGSGSYDAFDIW is shown. We found *IGHD3-10* to be the most likely D gene. There are 8 inserted bases in the junction. **(b)** The mean number of inserted bases is shown for several classes of cells occurring within the data, as well as the number of VDDJ instances. **(c)** For pairs of memory cells as in **Figure 1**, using varying CDRH3 amino acid identity, we show the light chain coherence (LCC) as a function of the number of inserted bases in the junction sequence. **(d)** Junctions for simulated heavy chains (from OLGA) and for naive cells in the data were aligned to the concatenated reference. Only those exhibiting no inserted bases were considered. The observed substitution rates are displayed. **(e)** CDRH3 lengths in amino acids are shown for naive and simulated junctions.

Our first new question was whether recurrent antibodies have fewer complex junctions than arbitrary antibodies. We analyzed each junction by finding the most likely D region (allowing for no D region, or a concatenation of two D regions to account for VDDJ junctions^27,28^), then aligned the antibody nucleotide sequence to the concatenation of the V(D)J reference sequences (**Figure 4a**). We considered the number of bases that were inserted in the antibody sequence (**Figure 4b** and **Extended Figure 6**). We found that recurrent antibodies have an order of magnitude fewer inserted bases, and that this is true for both naive and memory cells. This shows that recurrent antibodies have less complex junctions than arbitrary antibodies, and as expected, are more likely to recur by chance. In fact, most recurrent antibodies, whether naive or memory, have no inserted bases (**Extended Figure 6**). We analyzed this case in more detail for the case of naive cells (**Figure 4d**), finding that the junction substitution rate is lower by a factor of two in the recurrent case, again supporting our hypothesis that observed recurrent antibodies recur simply by chance.

We next asked if the observed recurrence rate was comparable to that expected by chance. Answering this question poses a quandary as it requires deep enough knowledge of recombination to accurately recapitulate the process by simulation. Thankfully other groups have tackled this challenging problem^29,30^. We generated random naive antibody sequences using the simulation program OLGA^29^, and compared the junction complexity and recurrence of simulated sequences to our data. We first asked how many recurrent naive antibodies are predicted by simulation, by making four groups of simulated antibodies of the same sizes as those as the groups of naive cells in our data, and then counting cross-donor recurrences of heavy chain gene and CDRH3 amino acid pairs. This yielded **78** cells (mean of ten replicates, {82, 76, 54, 81, 76, 69, 105, 84, 76, 77}), as compared to our observed value of **754** naive cells in our real data.

We next asked if any properties of the simulated naive antibodies could explain the underprediction of recurrences. We first noted that the number of inserted nucleotides for simulated sequences was **5.9**, as compared to **5.0** for naive cells (**Figure 4b**), suggesting that the simulation was roughly on track, while only partially explaining the recurrence discrepancy. We then turned to the substitution rate in the case where no junction insertions occurred (**Figure 4d**). Here we found a significant discrepancy: the mean substitution rate for simulated antibodies was **22%**, as compared to **16%** for real naive antibodies (range for the four donors: 16.21-16.61%). Clearly antibody sequences with fewer substitutions would be more likely to recur by chance. We also observed other discrepancies. First, the frequency of VDDJ junctions in the simulated antibodies was **17.8%**, as compared to **0.5%** (**Figure 4b**) in the real naive antibodies (and VDDJ antibodies would be far less likely to recur by chance). We note that CDRH3 lengths are shifted in a fashion consistent with the excess VDDJ recombinations (**Figure 4e**). Second, 8.8% of the simulated antibodies used the heavy chain V gene *IGHV3-NL1*, for which there appears to be no known full length sequence (beginning with a start codon) and which has only been observed in specific human populations^31,32^. Although collectively these data cannot fully explain the discrepancy between naive and simulated recurrence, they do suggest a possible reconciliation. Finally, we have investigated light chain coherence in antibodies that recur in four individuals. These are special antibodies, and therefore we wondered if the results would apply to all antibodies. While this cannot be answered using our data, we could ask if light chain coherence holds for the *relatively* more complex recurrent antibodies (**Figure 4c**). We find that recurrent antibodies with multiple inserted nucleotides in fact have higher light chain coherence than those with no inserted nucleotides. This lessened our concern that recurrent antibodies are particularly special. Indeed, our findings imply that all antibodies are recurrent, but at varying rates depending on their junction complexity and the prevalence of their cognate antigen. Our findings also suggest that except for their frequency, more complex antibodies do not behave differently with respect to light chain coherence.

## Discussion

Our work supports the following model, generalizing observed constraints on gene usage by some antibodies^2–8,10,12,14–24^. In nature, many heavy chain configurations yield effective binding of a given antibody target. However, for each of those, the cognate light chain is largely determined, at a rate of up to 80% at the level of light chain gene or paralog. We call this phenomenon *light chain coherence* and we observe it by looking for recurrences of heavy chains in memory and naive cells from four donors. The small number of donors biases our analysis towards junction regions with low complexity, though the same phenomenon is visible for more complex junctions that appear in our data. Our findings suggest that light chain coherence may apply to memory B cells in general.

While we generated V(D)J data for 1.6 million cells in this work, deeper data has been generated separately for heavy and light chains. This has enabled the identification of recurrences (public clonotypes) within such data, using strict definitions based on 100% CDRH3 amino acid identity^22,23^. We reinterpret these data here with some trepidation because of the differences in scale and technical approaches between the studies. We show here that previously described recurrences in these and other studies come from two types of B cells: naive B cells making “not yet functional” antibodies with minimal light chain coherence and memory B cells making functional antibodies with light chain coherence. The naive B cells’ lack of light chain coherence is expected given their lack of acquired functionality. Conversely, the memory B cells’ light chain coherence is expected because of their acquired functionality.

The simplest explanation for the recurrence we and others observe between naive cells is that their sequences repeat purely by chance. We show that in our data, recurrent naive cells (as well as memory cells) have markedly lower junction complexity, and it is thus no surprise that they recur. One might ask if the observed recurrence frequency is consistent with the mechanistic biology of V(D)J recombination. Answering that would require precise quantitative knowledge regarding this exquisitely complex process, and such knowledge does not exist. Moreover, we show that one simulator of this process does in fact predict recurrences, and although those recurrences are predicted to occur at a lower rate than we observe, we also show that the simulator generates significantly more complex sequences than those appearing in nature (**Figure 4**). With less complex sequences, simulated recurrence would be more frequent. Thus, we propose that naive sequences recur by chance. Conversely, recurrent memory sequences are a product both of chance and common exposure to related antigens. We suggest that recurrent naive sequences are instructive with respect to recombination and that recurrent memory sequences are instructive with respect to antibody function.

We postulate that light chain coherence implies functional coherence, and use this to suggest an amino acid substitution matrix, COSUM, appropriate for functional comparison and grouping of antibody sequences (**Extended Figure 5**). However, we do not claim to solve the problem of functional grouping. For this to be possible, at least two hurdles remain. First, all approaches (including ours) based on direct sequence comparison are naive to the structural consequences of amino acid changes. Rather than compare sequences, a more effective route to functional grouping may be to first computationally model antibody structures, from their sequences, and then compare those structures^33–37^. Second, far better truth data are needed in order to assess any method. It follows that while similar antibodies for the same antigen have been widely observed in multiple individuals, it is unknown how often antibodies to *different* antigens might be equally similar. Truth data targeted at such questions could comprise a suite of large datasets of naturally occurring antibodies, with one dataset for each of several antigens, along with binding data for each antibody. These data would be most powerful if they included nucleotide sequences (permitting *e*.*g*. consistent VDJ gene identification) and enough donor information to distinguish *bona fide* recurrence from clonal expansion within given individuals. Generation of such truth data at scale is feasible using existing methods^21,38–41^.

By virtue of how V(D)J recombination works, the light chain sequences of antibodies carry less information than the heavy chain sequences. Our work reveals that the light chain of functional antibodies is highly constrained, which implies that the light chains used in nature are the most functional ones: natural selection has won out. The complex dance between the heavy chain and the light chain is best studied at the level of individual cells—the context in which antibodies are produced, selected, and expanded. Although we do not yet understand why, we show that the choreography of this dance leads to a limited number of acceptable light chains. We suggest that antibody designers would be wise to actively look for optimal light chains used widely in nature, rather than focusing on the heavy chain alone. Similarly for bispecific antibodies, it could be advantageous to find two heavy chains whose native light chains are similar.

## Data Availability

All data are publicly available at https://figshare.com/articles/preprint/Functional_antibodies_exhibit_light_chain_coherence/19617633, including processed full-length V(D)J sequences and annotations. See **Methods** for informed consent and related information.

## Code Availability

All code to replicate key findings and figures of the paper are available at https://github.com/10XGenomics/enclone.

## Methods

For several details, we refer to ^42^, hereafter referred to as “enclone preprint”. We exhibit commands using executables in the enclone code and data of this work, at https://github.com/10XGenomics/enclone/tree/master/enclone_paper/which_code_does_what. A single line may be used to install the `enclone` executable and download the data, see [bit.ly/enclone](https://10xgenomics.github.io/enclone/). Use of the other executables requires compilation from source code at https://github.com/10XGenomics/enclone, and are in its directory `enclone_paper/src/bin`. We used version 0.5.175 of `enclone`. The `enclone` runs require about 145 GB memory and 20-40 minutes on a multi-core server (e.g. 24 cores). As an intermediate, we generated a file `per_cell_stuff`, with one line per cell, available [here](https://figshare.com/ndownloader/files/34846929) and facilitating direct analysis of the data of this work by other methods.

### Flow cytometry

We used a Sony MA900 cell sorter to purify single B cell suspensions from peripheral blood mononuclear cells from the four donors described in this paper. We used the following flow gating definitions for each population:

- <u>Naive</u>: live, CD3^−^, CD19^+^, IgD^+^, CD27^±^, CD38^±^, CD24^±^
- <u>Unswitched memory</u>: live, CD3^−^, CD19^+^, CD27^+^, IgD^low^, IgM^++^, CD38^±^, CD24^±^
- <u>Switched memory</u>: live, CD3^−^, CD19^+^, CD27^+^, IgD^−^, CD38^±^, CD24^±^, CD95^±^
- <u>Plasmablast</u>: live, CD3^−^, CD19^+^, CD27^+^, IgD^−^, CD38^++^, CD24^−^

The antibody panel we used was comprised of the following clones:

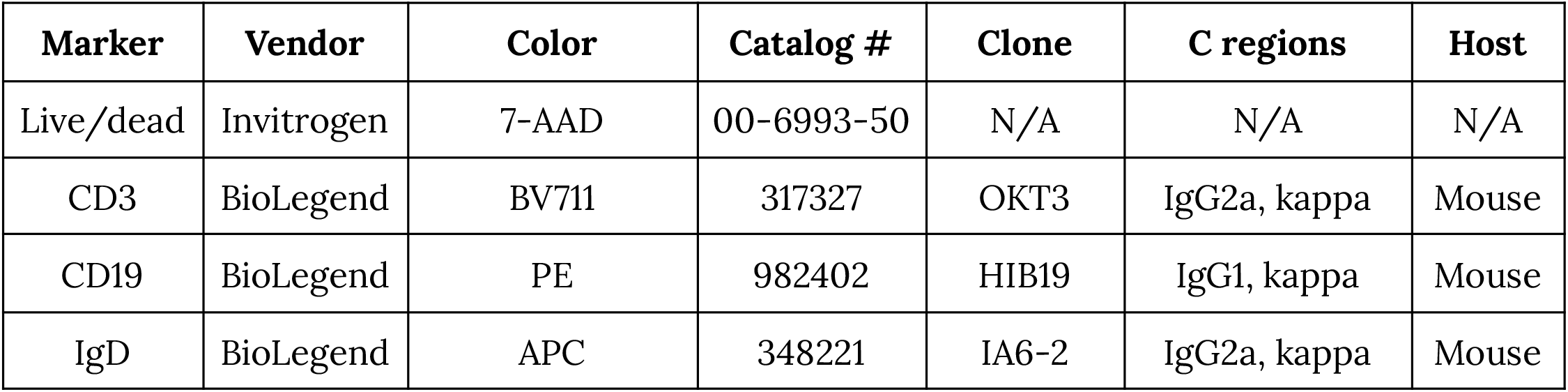
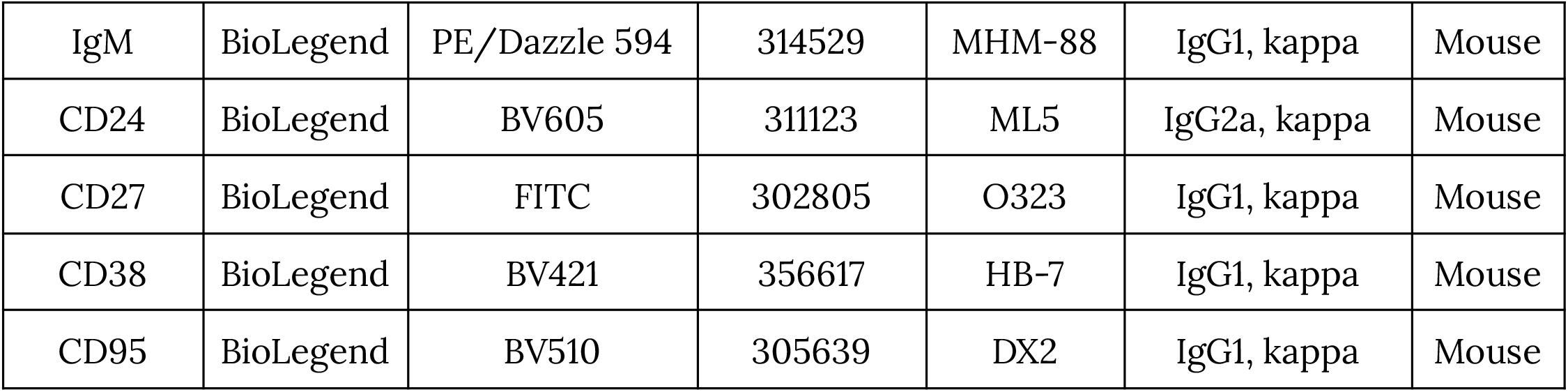

We titrated and developed the B cell fractionation panel using 20 million fresh PBMCs (AllCells, catalog # 3050363) from a healthy human donor whose cells were not used to generate single cell data in this paper. We thawed the cells per the 10x Genomics Demonstrated Protocol for Fresh Frozen Human Peripheral Blood Mononuclear Cells for Single Cell RNA Sequencing (CG00039, Revision D). Briefly, we resuspended cells in 20 μL of PBS / 2% FBS and incubated the cells on ice for 30 minutes in the dark. Before sorting, we washed the cells in 3x 1 mL of PBS / 2% FBS and then resuspended in 300 μL of PBS / 2% FBS for the sort step.

### Single cell data generation

Cells from four donors were flow sorted as naive, switched memory, unswitched memory, and plasmablast. V(D)J sequences were obtained using the 10x Genomics Immune Profiling Platform, using six Chromium X HT chips, and standard manufacturer methods. cDNA libraries were sequenced on the NovaSeq 6000 platform using several S4 flow cells. Certain memory B cell populations were relatively uncommon within certain donors. To account for this we spiked naive B cells into isolated memory cells from each donor to enable capture of a sufficiently large number of total B cells and unique sequences from each donor (see also **Extended Table 1**) targeting 20,000 cells recovered from each lane on the HT chip.

### Sequence simulation

We used OLGA v.1.2.4 to simulate 10 replicates of 1,408,939 heavy chain junctions. A reproducible Conda environment and scripts to generate these files (including random seeds) are provided at https://figshare.com/articles/preprint/Functional_antibodies_exhibit_light_chain_coherence/19617633. We used the default IGH model included with OLGA.

### Sequence logo plots

We used the ggseqlogo and msa R packages to align CDRH3 and CDRL3 sequences with the MUSCLE algorithm and to plot logo plots using position-wise entropy/Shannon information (y axis unit: bits). Letters were colored based on the properties of various amino acids. The amino acid-property color coding and other code necessary to reproduce these figures are publicly available as part of this paper.

### Light chain coherence optimized substitution matrix (COSUM)

We started with an amino acid substitution matrix having zeroes on the diagonal and ones elsewhere. We then iteratively and randomly perturbed the matrix by selecting at random two matrix entries, increasing one by a random amount and decreasing the other by another random amount. Values were truncated to one digit after the decimal point and capped at 8. The matrix was applied to memory cell pairs in the data, with each pair in a cell from a different donor, and sharing heavy chain V gene names and CDRH3 lengths. A perturbation was accepted if it increased the number of proximate pairs at 90% weighted percent identity and maintained light chain coherence of at least 75%. The calculation was stopped after 575,142 iterations (43 hours), at which point there were 7.8 times as many pairs.

### Allele inference

Donor alleles for V genes are partially inferred (as part of the enclone software, in the file `allele.rs`), using the following algorithm. The core concept is to pile up the observed sequences for a given V gene and identify variant bases. If we used all cells (one sequence per cell), then the sequences would be biased by clonal expansion and thus yield incorrect alleles. Ideally we would instead use just one cell per clonotype that uses the given V gene. However the order of operations is that we first compute donor alleles, and then compute clonotypes. Therefore we use a heuristic for picking cells that does not depend on knowing the clonotypes. The heuristic is that we pick just one cell among those using the given V gene, and that share the same CDRH3 length, CDRL3 length, and partner chain V and J genes. The pileup is then made from the V gene sequences of these cells.

Next, for each position along the V gene, excluding the last 15 bases (to avoid the junction region), we determine the distribution of bases that occur within these selected cells. We only consider those positions where a non-reference base occurs at least four times and represents at least 25% of the total. Then each cell has a footprint relative to these positions; we require that these footprints satisfy similar evidence criteria. Each such non-reference footprint then defines an “alternate allele”. We do not restrict the number of alternate alleles because they could arise from duplicated gene copies. The ability of the algorithm to reconstruct alleles is limited by the depth of coverage (counted in “non-redundant” cells) of a given V gene. Moreover the algorithm cannot identify germline mutations which occur in the terminal bases of the V gene, inside the junction region.

## Acknowledgements

We wish to express our gratitude to the donors for their patience in undergoing apheresis and donating their blood. We thank Pat Marks, Mike Stubbington, Sarah Taylor, Preyas Shah and Vijay Kumar for their comments and suggestions.

## Author contributions

D.B.J. and W.J.M. were responsible for conceptualization, formal analysis, methodology, software, supervision, validation, and writing of the original draft manuscript. D.B.J., P.S., B.A.A, N.R., D.S.R., and W.J.M. were responsible for data curation. All authors were responsible for investigation. D.B.J., P.S., D.S.R., and W.J.M. were responsible for project administration. D.B.J., P.S., N.R., and W.J.M. were responsible for visualization. All authors were responsible for review and editing of the final draft manuscript. Funding and resources were provided by 10x Genomics, Inc.

## Extended Data Tables and Legends

**Extended Table 1.**
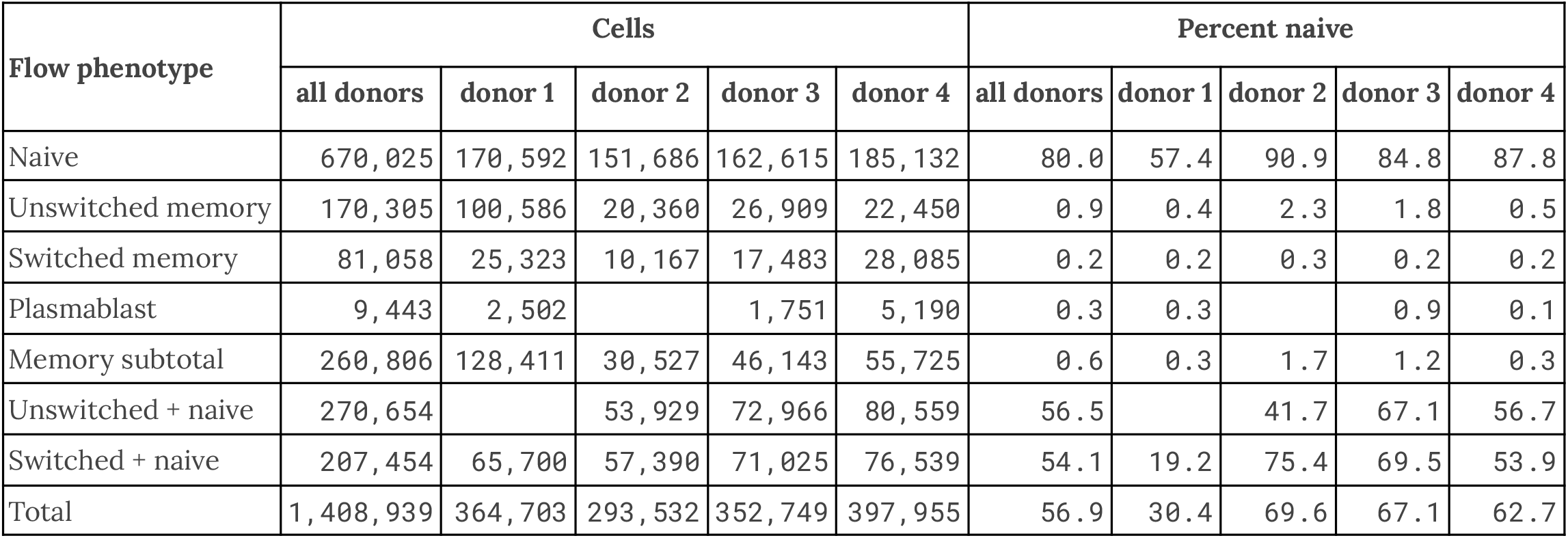
Flow phenotype categories by donor with total number of cells and relative naive fraction. Total numbers of cells captured via fluorescence-activated flow cytometry with exactly two chains are shown, along with the fraction of naive (d_*ref*_ = 0) sequences. Cells were sorted for naive, unswitched, switched and plasmablast, and in some libraries, sort categories were combined. Entries are blank if no data were generated. The table only accounts for cells that exhibited exactly one heavy and one light chain, and which were determined to lie in a valid clonotype having exactly two chains.

| Flow phenotype | Cells |  |  |  |  | Percent naive |  |  |  |  |
| --- | --- | --- | --- | --- | --- | --- | --- | --- | --- | --- |
|  | all donors | donor 1 | donor 2 | donor 3 | donor 4 | all donors | donor 1 | donor 2 | donor 3 | donor 4 |
| Naive | 670,025 | 170,592 | 151,686 | 162,615 | 185,132 | 80.0 | 57.4 | 90.9 | 84.8 | 87.8 |
| Unswitched memory | 170,305 | 100,586 | 20,360 | 26,909 | 22,450 | 0.9 | 0.4 | 2.3 | 1.8 | 0.5 |
| Switched memory | 81,058 | 25,323 | 10,167 | 17,483 | 28,085 | 0.2 | 0.2 | 0.3 | 0.2 | 0.2 |
| Plasmablast | 9,443 | 2,502 |  | 1,751 | 5,190 | 0.3 | 0.3 |  | 0.9 | 0.1 |
| Memory subtotal | 260,806 | 128,411 | 30,527 | 46,143 | 55,725 | 0.6 | 0.3 | 1.7 | 1.2 | 0.3 |
| Unswitched + naive | 270,654 |  | 53,929 | 72,966 | 80,559 | 56.5 |  | 41.7 | 67.1 | 56.7 |
| Switched + naive | 207,454 | 65,700 | 57,390 | 71,025 | 76,539 | 54.1 | 19.2 | 75.4 | 69.5 | 53.9 |
| Total | 1,408,939 | 364,703 | 293,532 | 352,749 | 397,955 | 56.9 | 30.4 | 69.6 | 67.1 | 62.7 |

**Extended Table 2.**
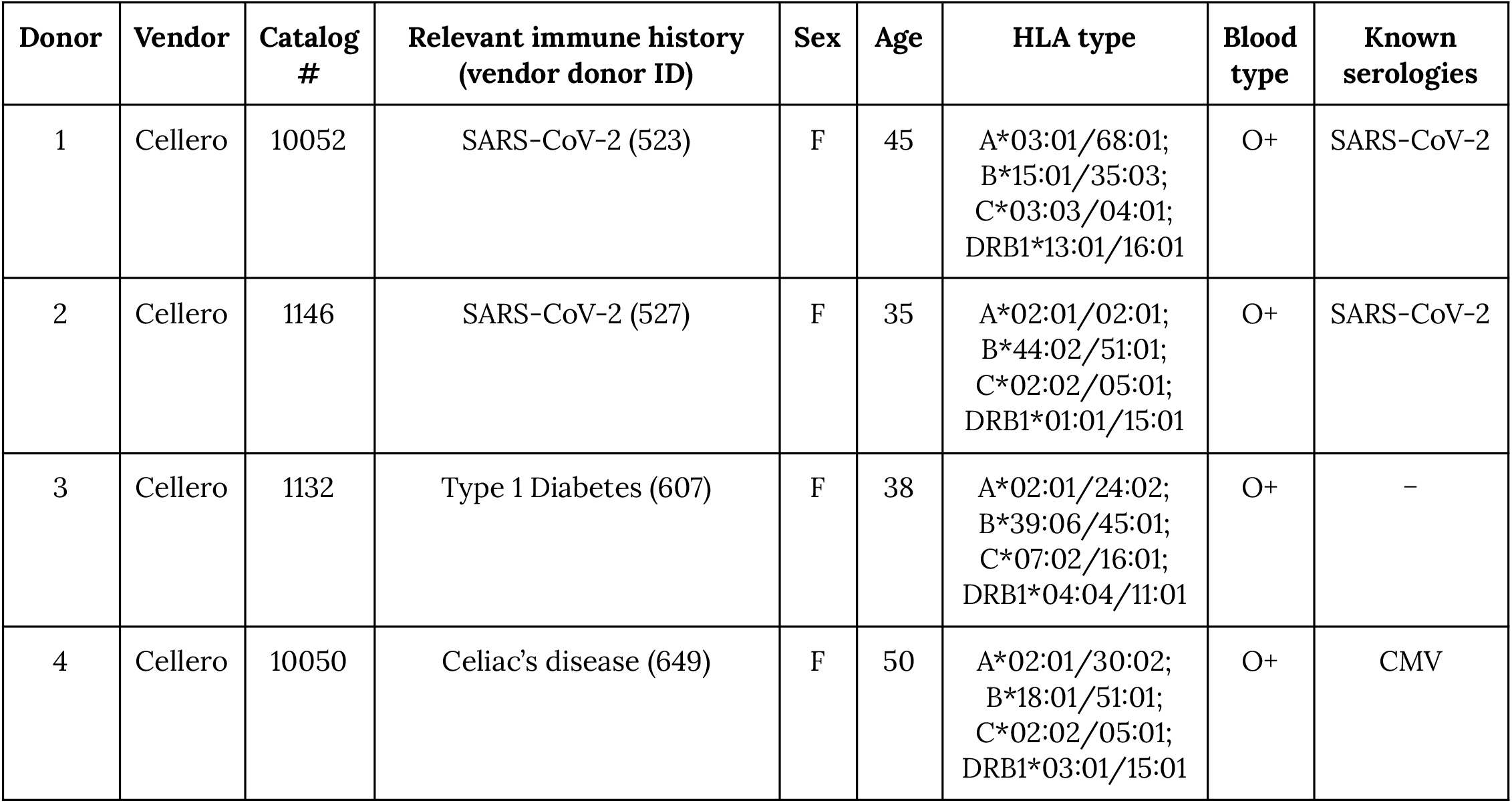
Donor information. All donors were appropriately consented for genomic data use and release under protocols reviewed by independent IRB boards consulted by the vendor. Samples from the donors were tested by the vendor and confirmed to be both seronegative and not detectably infected with HIV-1, HIV-2, hepatitis B, hepatitis C, or HTLV-1. HLA typing, serology, and blood typing were also performed by the vendor. <u>Donor 523 clinical timeline</u>. Donor 523 tested positive for COVID-19 via RT-PCR of nasopharyngeal swabs on day 0 and was hospitalized from day -5 to day 0. She tested negative for COVID-19 at day 18, and had a plaque reduction neutralization test titer of 1:>2560 at day 44. She donated plasma and cells on day 65. <u>Donor 527 clinical timeline</u>. Donor 527 tested positive for COVID-19 via RT-PCR of nasopharyngeal swabs on day 0 and was not hospitalized. She tested negative for COVID-19 at day 15, and had a plaque reduction neutralization test titer of 1:20 at day 57. She donated plasma and cells on day 75.

**Extended Figure 1.**
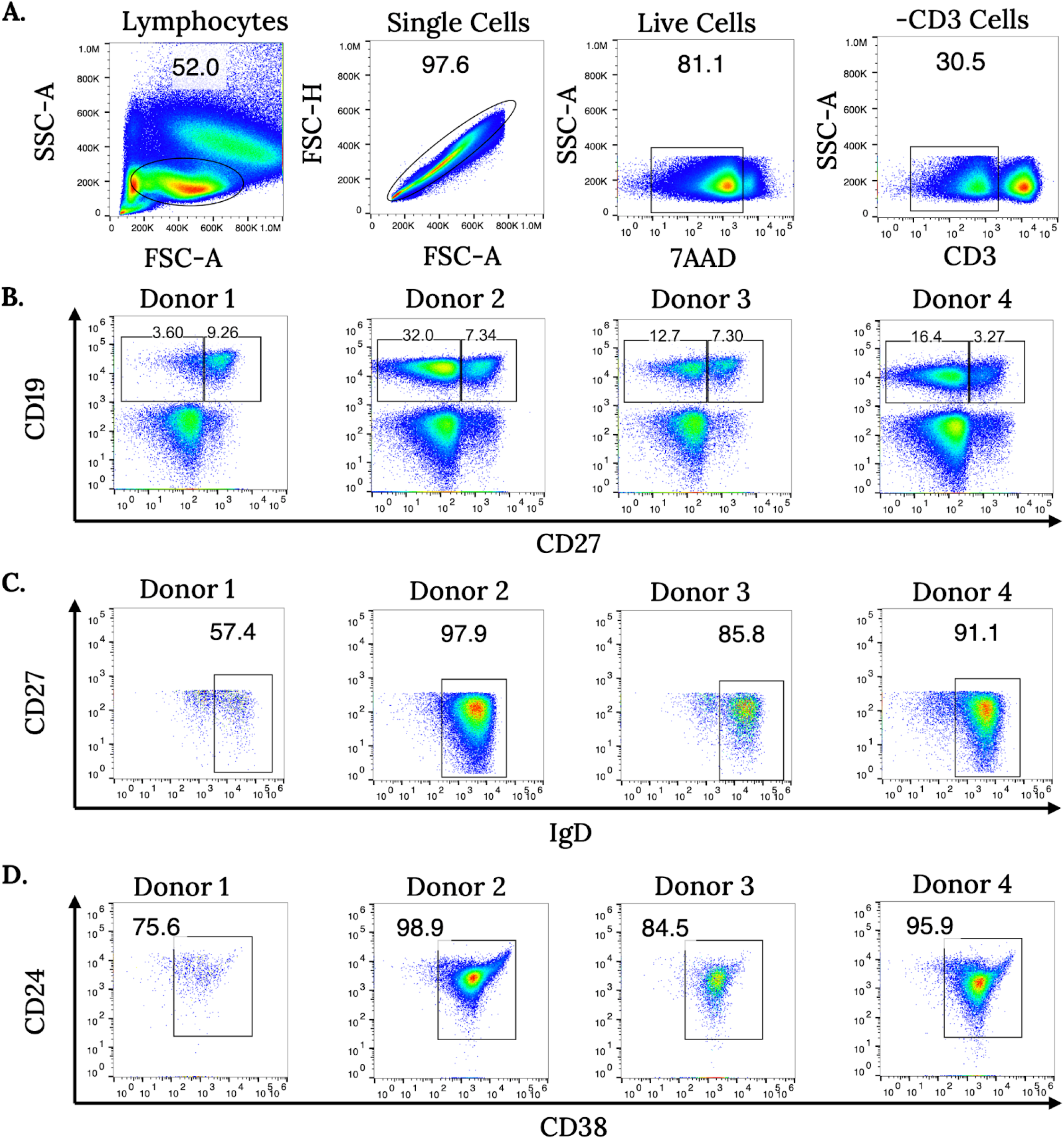
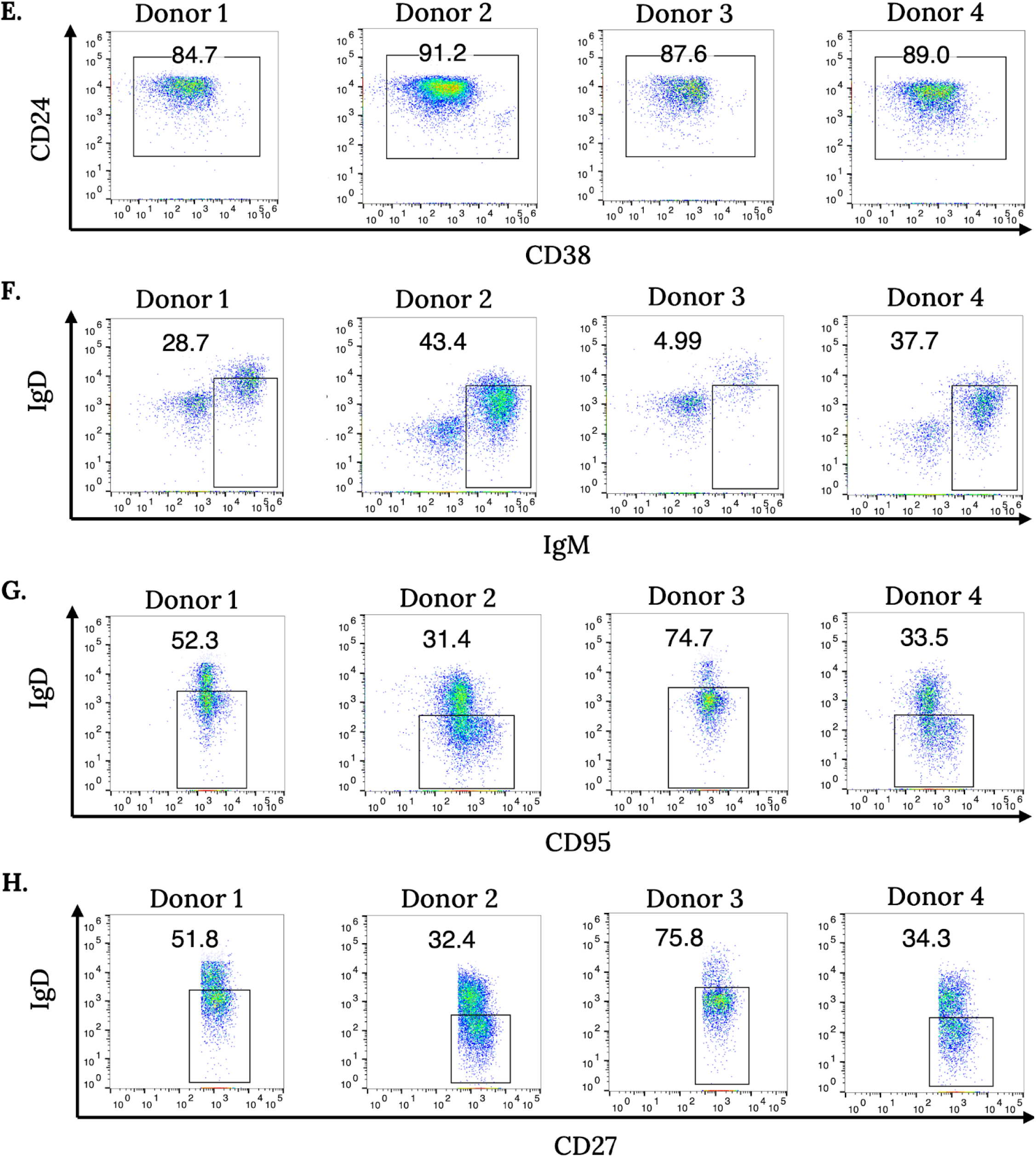
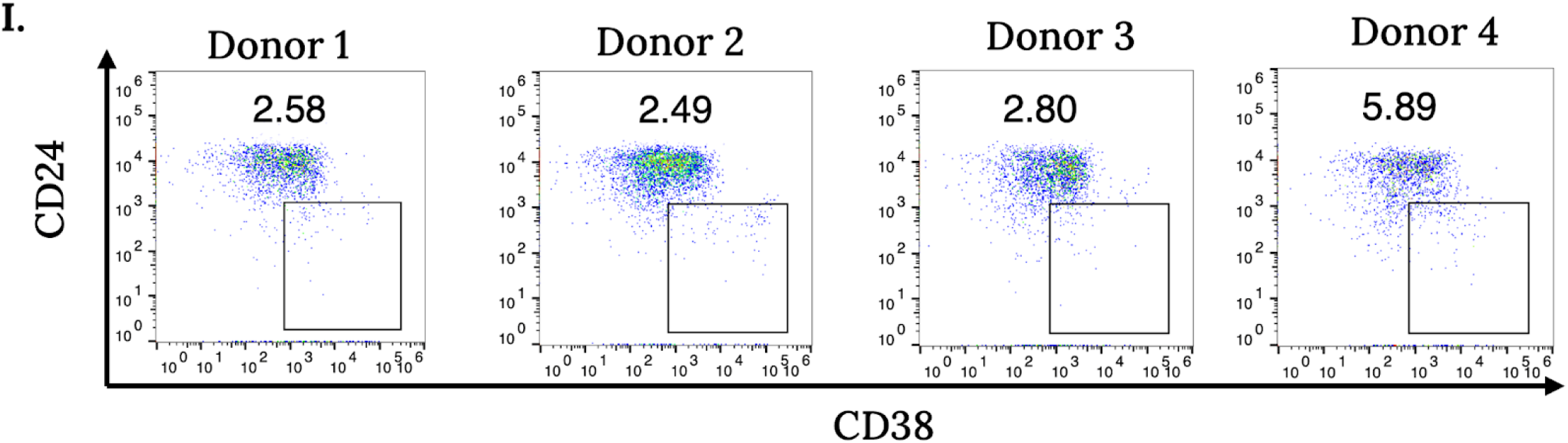
Flow cytometry gating schemes for B cell subsets. Gating strategy for isolating naïve B cells. Panels (a-d) show naive cell gating, panels (e-g) show memory cell gating, and panels (h-i) show plasmablast gating. **(a)** Hierarchical gating scheme for lymphocytes, single cells, live cells, and CD3-negative cells. **(b)** We gated CD19+CD27± cells from CD3− cells for further analysis. Donor samples displayed noticeable differences in CD19 and CD27 expression. **(c)** We analyzed CD19+CD27− cells for surface IgD expression and gated IgD+ cells for further analysis. **(d)** We selected naïve B cells by sorting CD19+CD27–IgD+CD24±CD38± B cells. **(e)** For memory cell gating, we selected CD19+CD27+ cells from (b) for CD24+CD38+ positivity. **(f)** We analyzed cells from (e) and isolated unswitched memory cells using IgD±IgM++ gating. **(g)** We analyzed cells from (e) and isolated switched memory cells using IgD−CD95+ gating. **(h)** We analyzed CD19+CD27+ cells from (b) and gated the IgD-CD27+ population. **(i)** We sorted plasmablasts using CD24–CD38++ gating.

**Extended Figure 2.**
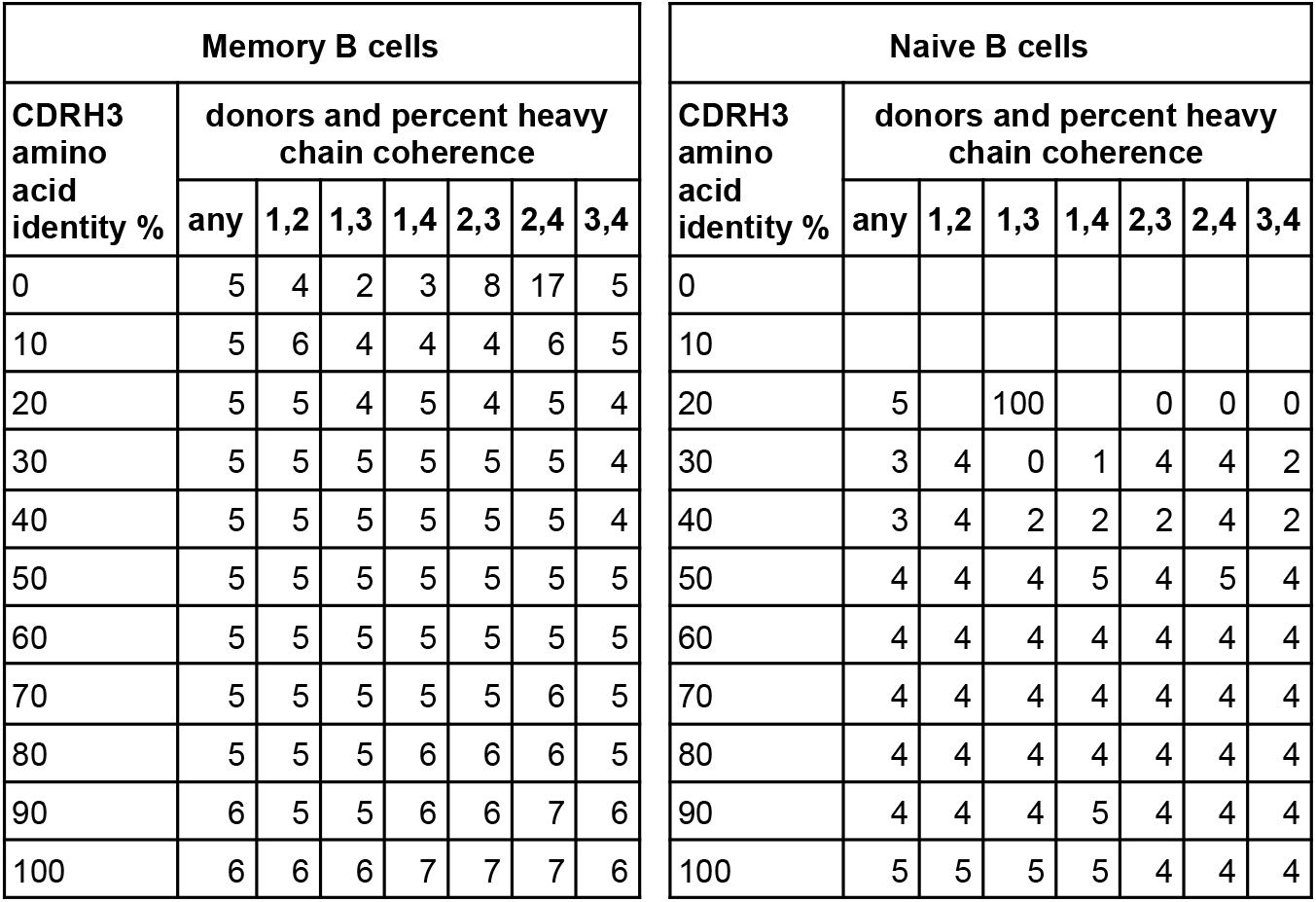
Light chain coherence does not imply heavy chain coherence. Data are shown as in **Figure 1**, except that the role of heavy and light chains is reversed, and paralogs are not considered. Entries are blank in cases where there was no data because no heavy chain sequences could be compared at a given CDRH3 percent identity threshold. A value of 0 represents 0% heavy chain concordance.

**Extended Figure 3.**
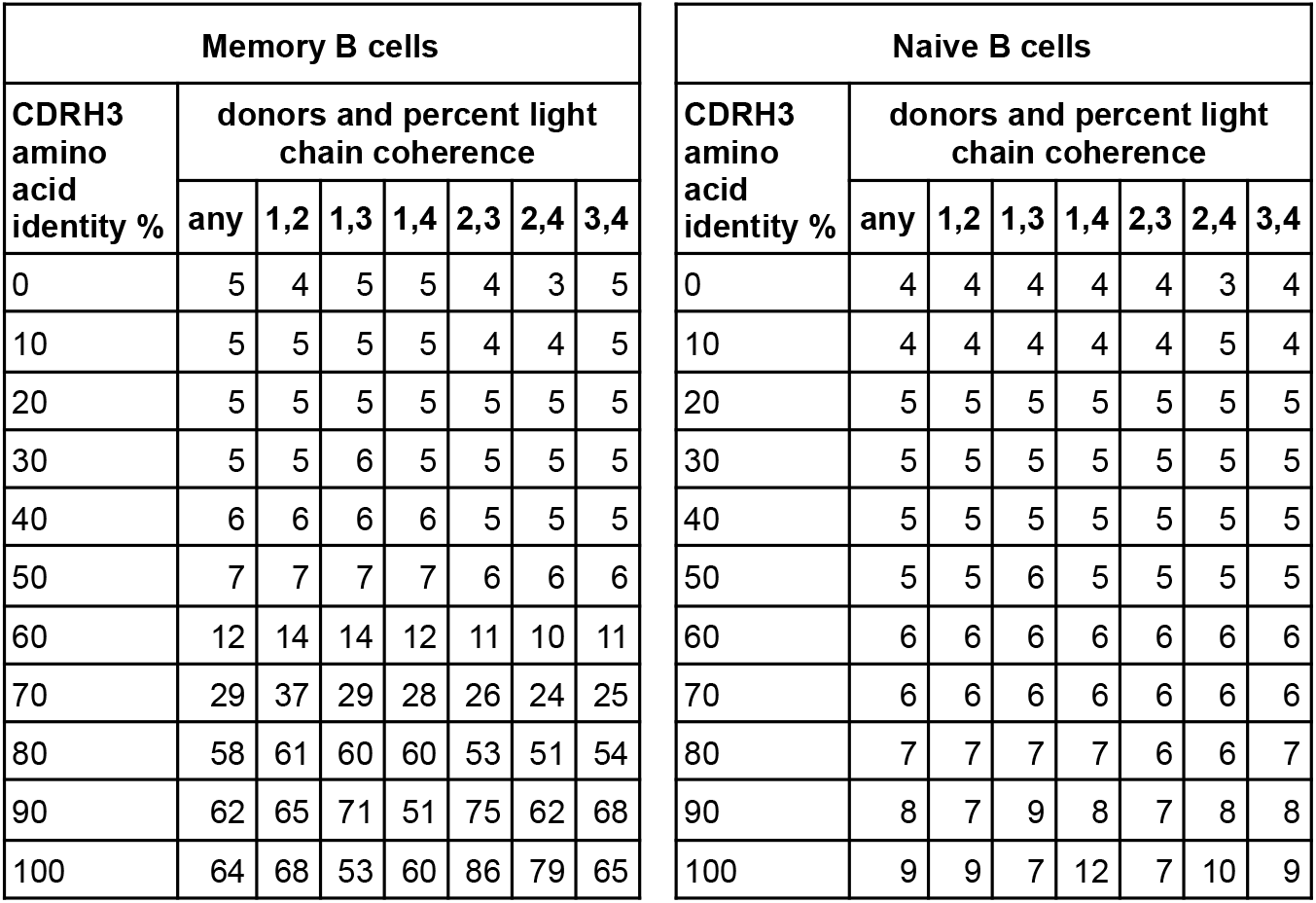
Light chain coherence in memory B cells (public antibodies), without identifying light chain V gene paralogs. Data are shown as in **Figure 1**, except that light chain V gene paralogs are not treated as the same.

**Extended Figure 4.**
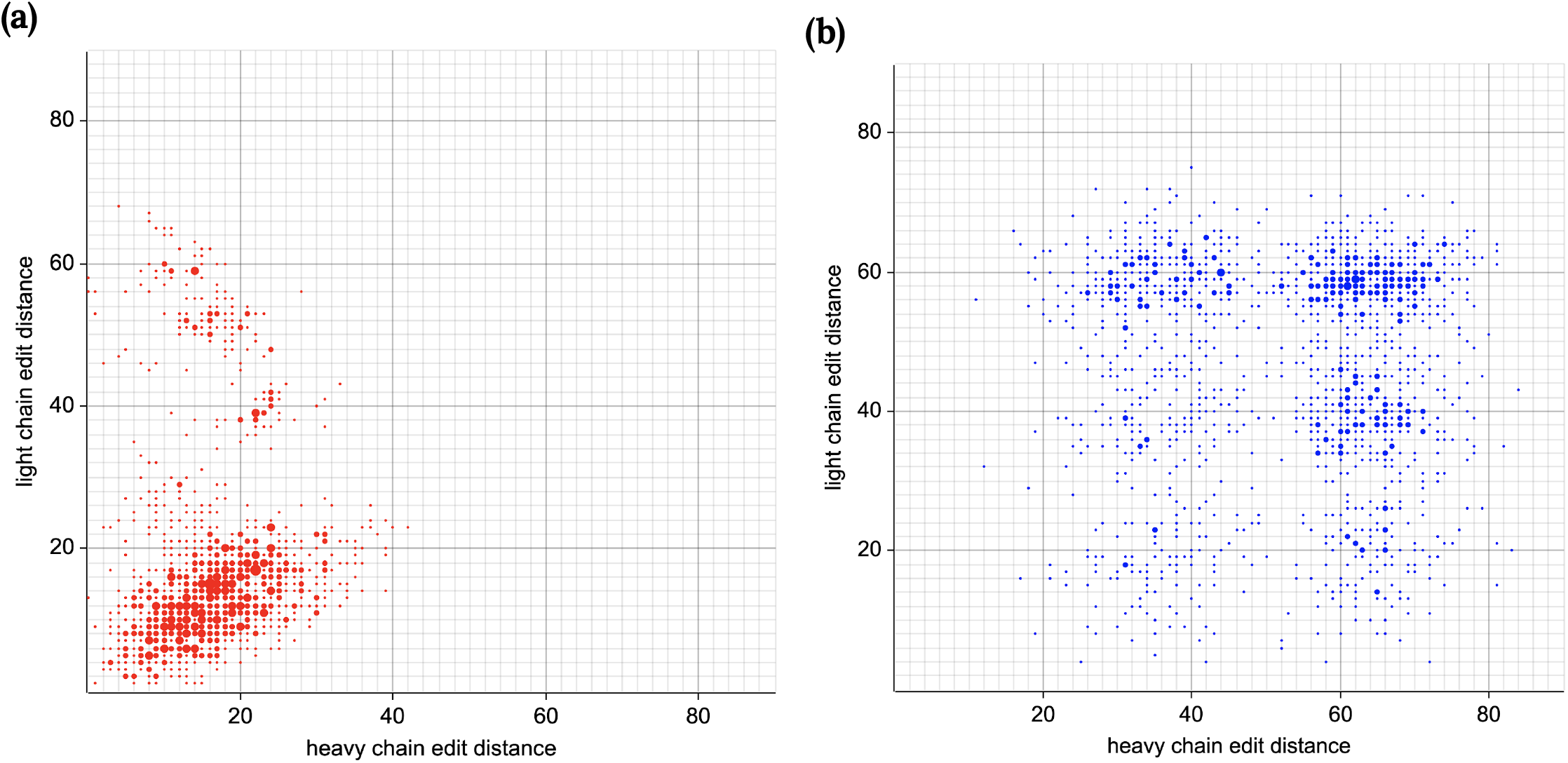
Light chain coherence is visible by sequence similarity. Each point represents a pair of memory cells from different donors. Heavy and light chain edit distances are plotted, using the amino acids starting at the end of the leader and continuing through the last amino acid in the J segment. Points with identical coordinates are combined by showing a large point whose area is proportional to the number of such points. **(a)** Cell pairs are displayed if the two cells in the pair have the same CDRH3 amino acid sequence. To increase readability, only one third of such pairs were selected at random for display. Of the pairs, **78%** have light chain edit distance ≤ 20. **(b)** [control] The same number of cell pairs were selected at random for display, without regard to CDRH3. Of the pairs, **9%** have light chain edit distance ≤ 20.

**Extended Figure 5.**
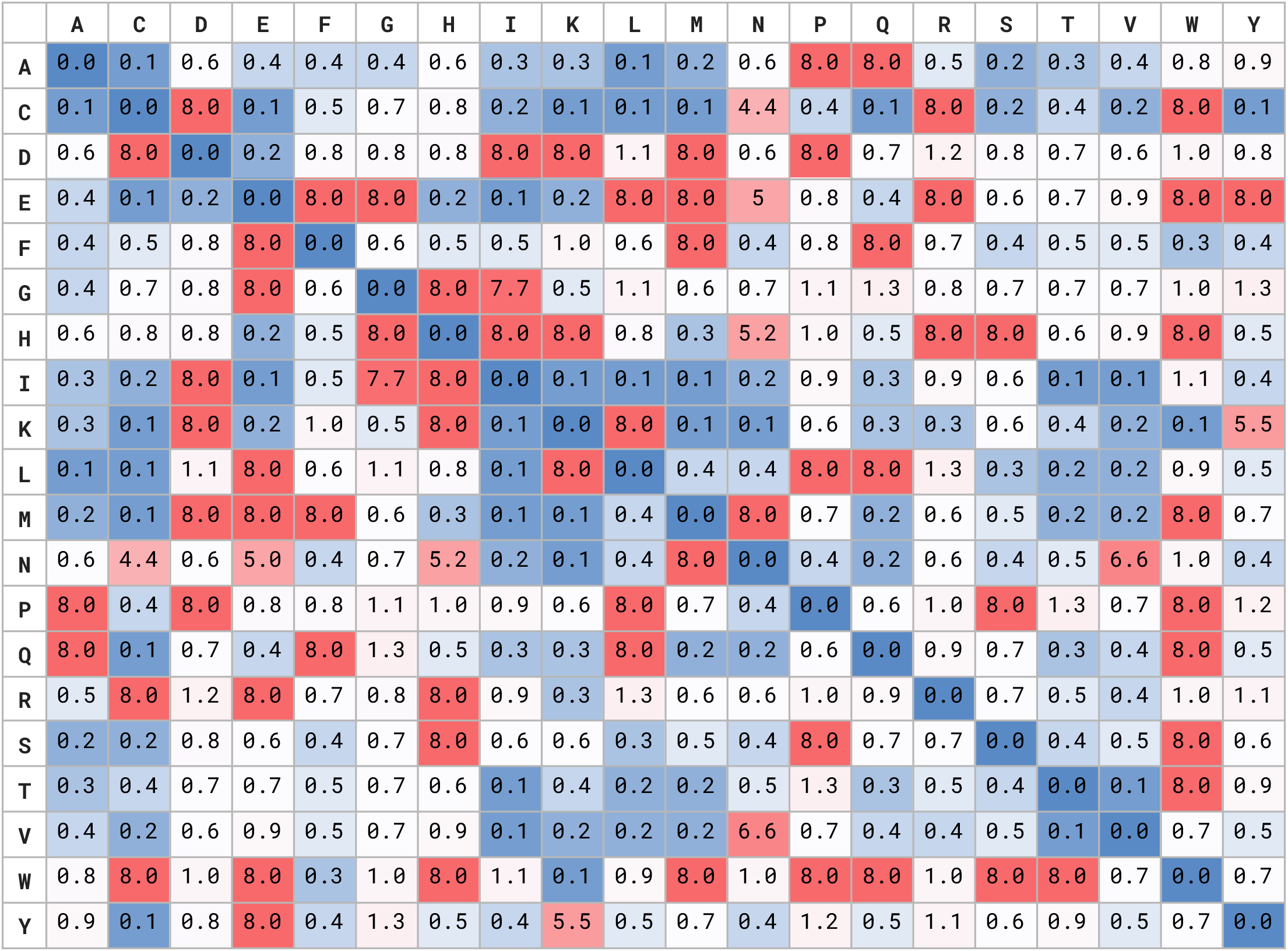
Light chain coherence optimized substitution matrix (COSUM). We found an amino acid substitution matrix M relative to which the CDRH3 sequences for many cell pairs in the data are close, and for which those cell pairs have at least 75% light chain coherence (**Methods**).

**Extended Figure 6.**
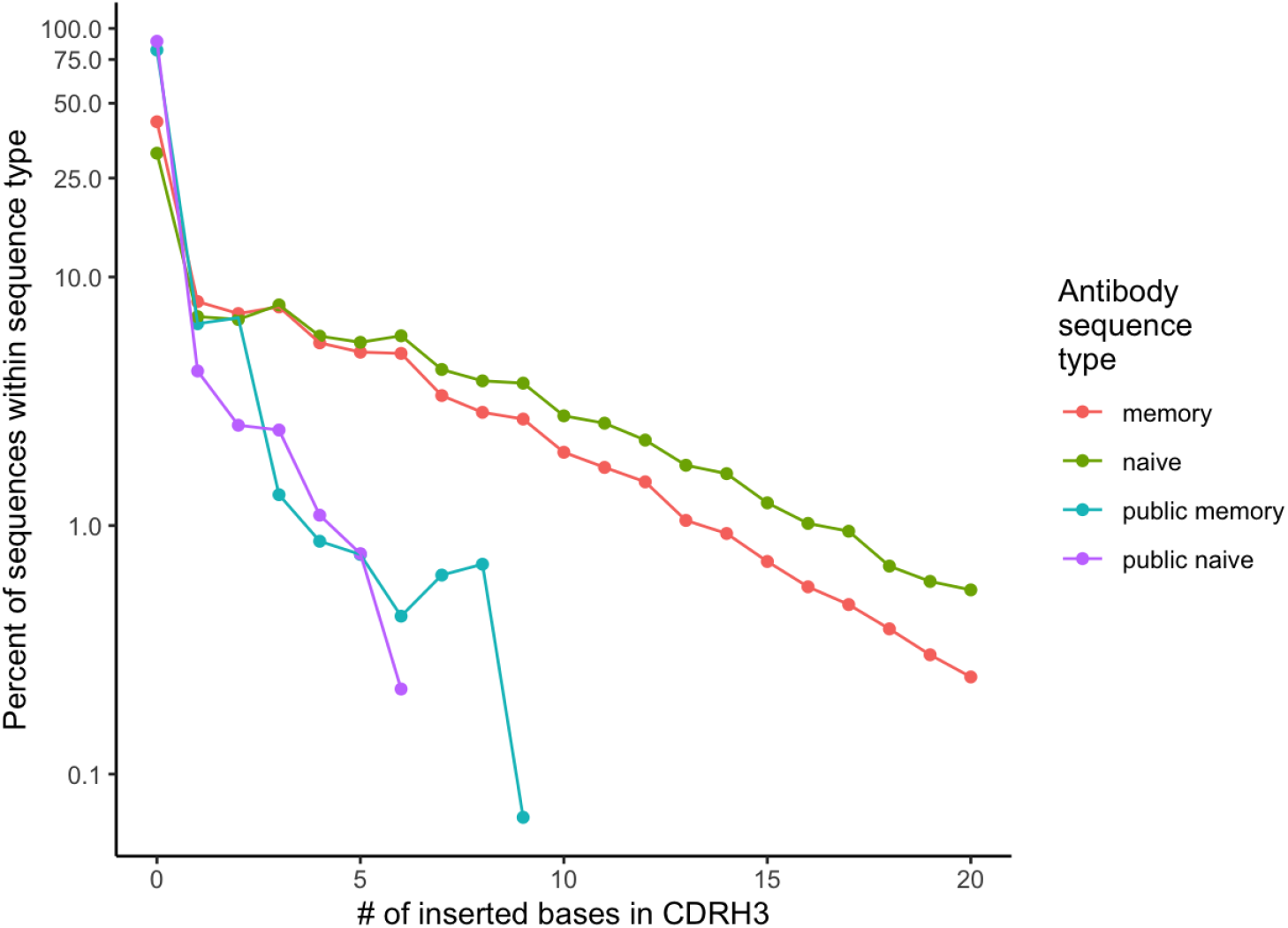
Inserted bases in CDRH3 for types of antibodies. For each of four types of antibodies in the data, we computed the number of inserted bases in the heavy chain junction region, relative to the concatenated VDJ (or in some cases VJ or VDDJ) reference sequence. Most of the inserted bases are N1 or N2 insertions. The frequency is shown as a function of the number of inserted bases.

